# Design of Antigen-Specific Antibody CDRH3 Sequences Using AI and Germline-Based Templates

**DOI:** 10.1101/2024.03.22.586241

**Authors:** Toma M. Marinov, Alexandra A. Abu-Shmais, Alexis K. Janke, Ivelin S. Georgiev

## Abstract

Antibody-antigen specificity is engendered and refined through a number of complex B cell processes, including germline gene recombination and somatic hypermutation. Here, we present an AI-based technology for de novo generation of antigen-specific antibody CDRH3 sequences using germline-based templates, and validate this technology through the generation of antibodies against SARS-CoV-2. AI-based processes that mimic the outcome, but bypass the complexity of natural antibody generation, can be efficient and effective alternatives to traditional experimental approaches for antibody discovery.

---

Monoclonal antibodies are an effective therapeutic modality against a wide range of diseases, including infectious disease, cancer, autoimmunity, and others ^1, 2^. The traditional antibody discovery process requires access to source samples with previous exposure to the antigen target, followed by sifting through an enormous pool of potential antibody candidates to pick leads that subsequently must be validated for specificity and function toward the target antigen ^3^. As a result, antibody lead discovery is typically associated with significant logistical challenges and high levels of inefficiency, and more effective and efficient approaches are critically needed ^4, 5^.

With recent advances in artificial intelligence (AI) and computational biology, the feasibility and accuracy of modeling protein sequence and structure has increased immensely, with effective applications to drug, vaccine, and biosensor design, among many others ^6, 7^. In the realm of antibodies, optimization of existing antibody candidates through targeted redesign of the initial sequence/structure of a traditionally sourced antibody has been accomplished ^8, 9^. However, it remains an open question whether such computationally driven approaches can be utilized to design human antibodies de novo, rather than redesign an existing antibody.

Here, we present a method for AI-based de novo design of antigen-specific antibody CDRH3 sequences that aims to mimic the outcome, but bypass the complexity, of natural antibody generation against a specific antigen target. Monoclonal antibodies are the secreted form of the B cell receptor on B cells, which in humans consists of a pairing of a heavy chain (HC) and a light chain (LC) protein. HC-LC pairing is one of the mechanisms that enables diversification of the antibody repertoire, along with germline gene recombination and somatic hypermutation in each of the HC and LC. Germline gene recombination in the HC involves three gene segments, V (variable), D (diversity) and J (joining), with the recombined HC retaining the majority of the V and J genes, while the segment in-between forms a hypervariable loop called the complementarity determining region 3 (CDRH3). The process in the LC is similar but only V and J genes recombine, resulting in less (though still notable) diversity in the CDRL3 region. In addition to significant contributions by the two CDR3 regions, antigen recognition by antibodies is achieved through critical interactions with parts of the HC and LC V-genes (VH and VL, respectively). Conceptually, nature uses a discrete set of building blocks and generally stochastic processes to assemble and modify these building blocks, in order to generate a diverse repertoire of antibodies that can tackle the immense diversity of potential antigen targets. Despite this intrinsic diversity of human antibody repertoires, the phenomenon of “public” antibody clonotypes that are found in multiple individuals and that utilize the same VH that exhibits conserved critical interactions with cognate antigen, have been identified for a number of antigens, including the SARS-CoV-2 spike ^10, 11^. The existence of such public antibodies suggests that there are reproducible, and generalizable, rules of antibody-antigen interactions.

Machine learning approaches are particularly suited to identifying associations between inputs and outcomes, without necessarily explicitly deciphering the complex underlying relationships. Recently, large language models (LLM), which have been successfully applied to numerous areas outside of language processing ^7, 12^, have been shown to effectively capture the fundamental rules of protein sequence and function by viewing proteins as a string (a sequence of characters). Such models have also recently been adapted to generate novel antibody sequences ^13, 14^, but the question remains whether these AI-generated antibodies can successfully target an antigen of interest, thus providing a novel and efficient method for the discovery of antigen-specific antibodies.

To address this question, we set out to explore whether an AI-based pipeline for antibody discovery can successfully generate antibody sequences against a target antigen. To validate this approach, we chose to design antibodies against a specific epitope on SARS-CoV-2 spike that is known to be targeted by a public class of antibodies utilizing the VH3-53 (or the highly similar VH3-66) germline gene ^11^. We then set up the computational problem to interrogate whether, given a public germline gene template and a specific structural epitope target, we can de novo design and down-select CDRH3 sequences with specificity for the target antigen. If successful, such an approach would mimic the outcome of natural antibody generation observed at the population level (hence, the choice of a public class of antibodies).

To that end, we first utilized the IgLM language model ^13^ to generate 1,000 de novo CDRH3 sequences flanked by germline V and germline J. The generated CDRH3 regions had substantial levels of diversity in sequence composition (**Figure 1A**) and length (**Figure 1B**) and were generally distinct from known CDRH3 sequences for natural SARS-CoV-2 antibodies from the CoV-AbDab database (**Figure 1C**). We then modeled the structure of all 1,000 generated antibody HC (including the generated CDRH3 and flanking germline V/J) using ImmuneBuilder^15^. When compared to the structure of a known antibody from this class (BD-604^16^), the generated CDRH3s exhibited a range of structural similarity levels (**Figure 1D**).

**Figure 1 Caption:**
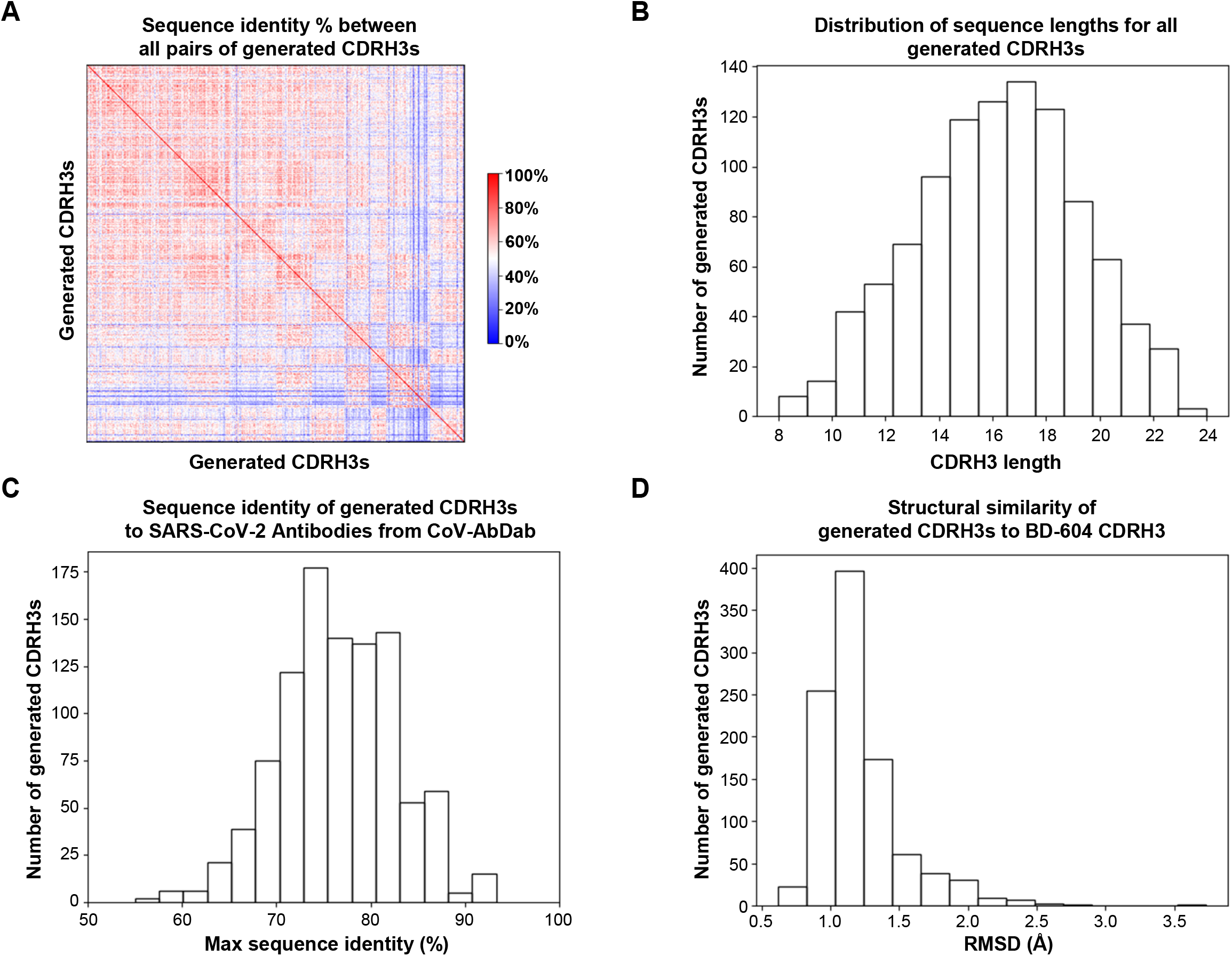
Characteristics of AI-generated antibody CDRH3 sequences. (A) The percent sequence identity between all pairs of generated CDRH3 sequences (rows/columns) is shown as a heatmap (blue: 0%, white: 50%, red: 100%). (B) The distribution of sequence lengths (x-axis) for the generated CDRH3s is shown with numbers of CDRH3s for each length group shown by bar height on the y-axis. (C) For each generated CDRH3 sequence, the maximum sequence identity (x-axis) to CDRH3 sequences for antibodies from the CoV-AbDab database was computed, and the distribution is shown as a histogram. (D) For each generated CDRH3, the distribution of the RMSD (x-axis) of the modeled structure when compared to the known structure of the CDRH3 of the SARS-CoV-2 antibody BD-604 is shown as a histogram.

To better understand the ability of these computational methods to generate antibody sequences with target antigen specificity, we selected 14 antibody candidates for experimental validation. To increase the likelihood of identifying positive hits, we focused on antibodies that: (a) were mostly associated with high predicted structural CDRH3 similarity to BD-604, and (b) were different in sequence to known VH3-53/VH3-66 SARS-CoV-2 antibodies (**Figure 2**). Recombinant monoclonal antibodies were tested for binding to spike protein by enzyme-linked immunosorbent assay (ELISA). As an initial screen, the 14 antibodies (formed by pairing the designed heavy chain with the BD-604 light chain) were tested for binding to the SARS-CoV-2 index variant (**Figure 3A**). Two of the antibodies (36 and 843) showed signal above background at lower antibody concentrations, and were therefore selected for further experimental characterization. Notably, the structural models for both of these antibodies had <1Å CDRH3 RMSD compared to BD-604 (**Figure 2**). We next tested binding of these two antibodies against SARS-CoV-2 index spike, alongside the BD-604 antibody. In addition, we also tested versions of antibodies 36 and 843 with the mature BD-604 VH that includes 5 somatic hypermutation changes compared to germline; we will refer to these antibodies as gv36, gv843 for the germline VH version and mv36, mv843 for the mature VH version. Antibodies gv36 and gv843 showed saturation binding to spike, although to a lesser extent compared to the BD-604, mv36, and mv843, which all showed similar levels of antigen binding (**Figure 3B**). These binding patterns were retained when the antibodies were tested against a variety of different variants of the SARS-CoV-2 virus ^17^, including XBB.1 and BQ.1.1, with the only exception of BA.1 (which had occurred earlier in the pandemic compared to the more recent variants XBB.1 and BQ.1.1) where only BD-604 showed binding (**Figure 3C**).

**Figure 2 Caption:**
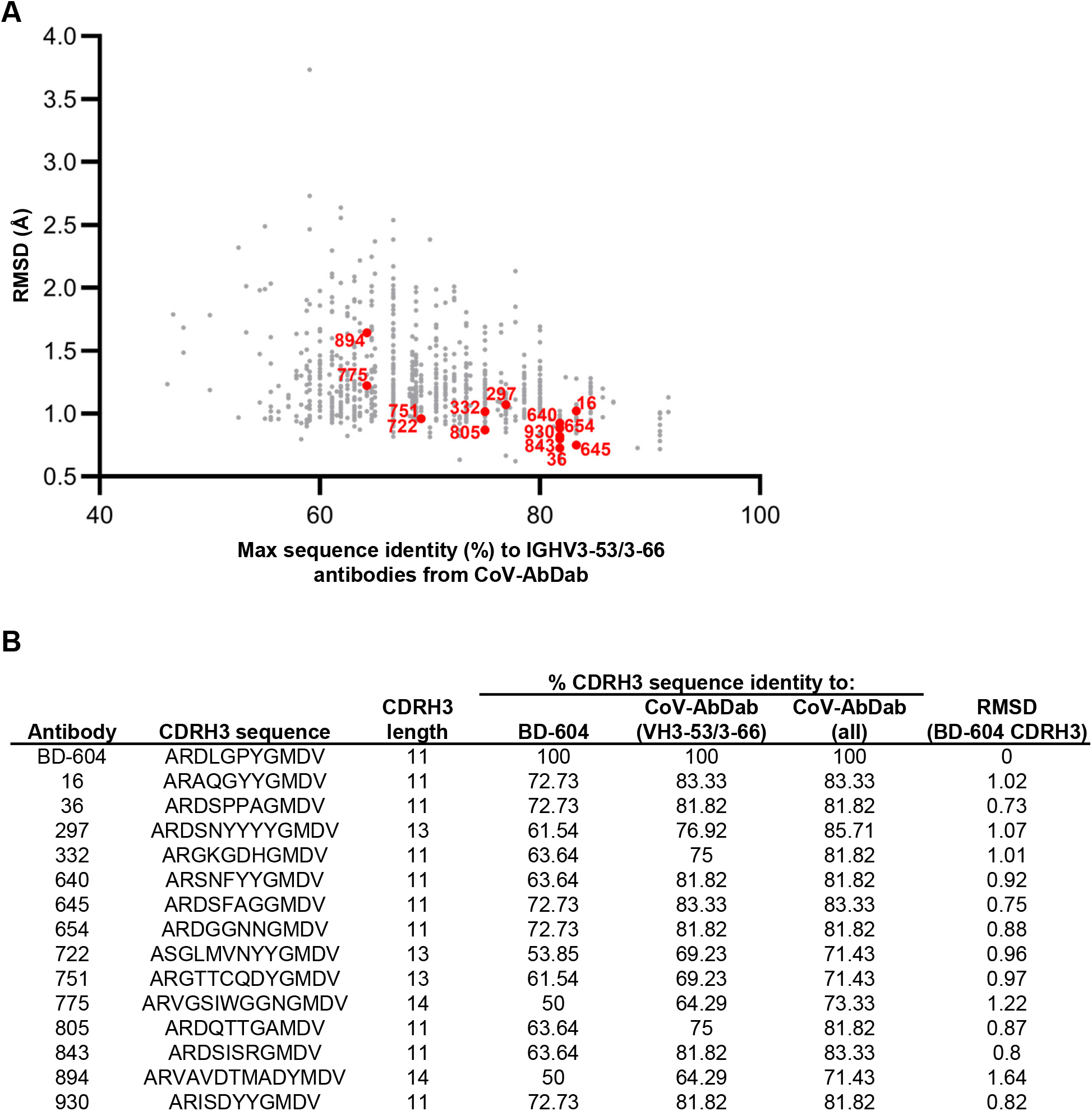
Characteristics of antibodies selected for validation. (A) For each generated CDRH3 (dots), shown are the percent sequence identity to CDRH3s for antibodies using the public IGHV3-53/3-66 germline gene from the CoV-AbDab database (x-axis) vs. RMSD to BD-604 (y-axis). The 14 antibodies that were selected for experimental validation are highlighted in red. (B) For each of BD-604 and the 14 selected AI-generated antibodies (rows), shown are the CDRH3 sequence and length; the % CDRH3 sequence identity to BD-604, antibodies using the public IGHV3-53/3-66 germline gene from the CoV-AbDab database, and all antibodies from the CoV-AbDab database; and RMSD (in Å) to the BD-604 CDRH3.

**Figure 3 Caption:**
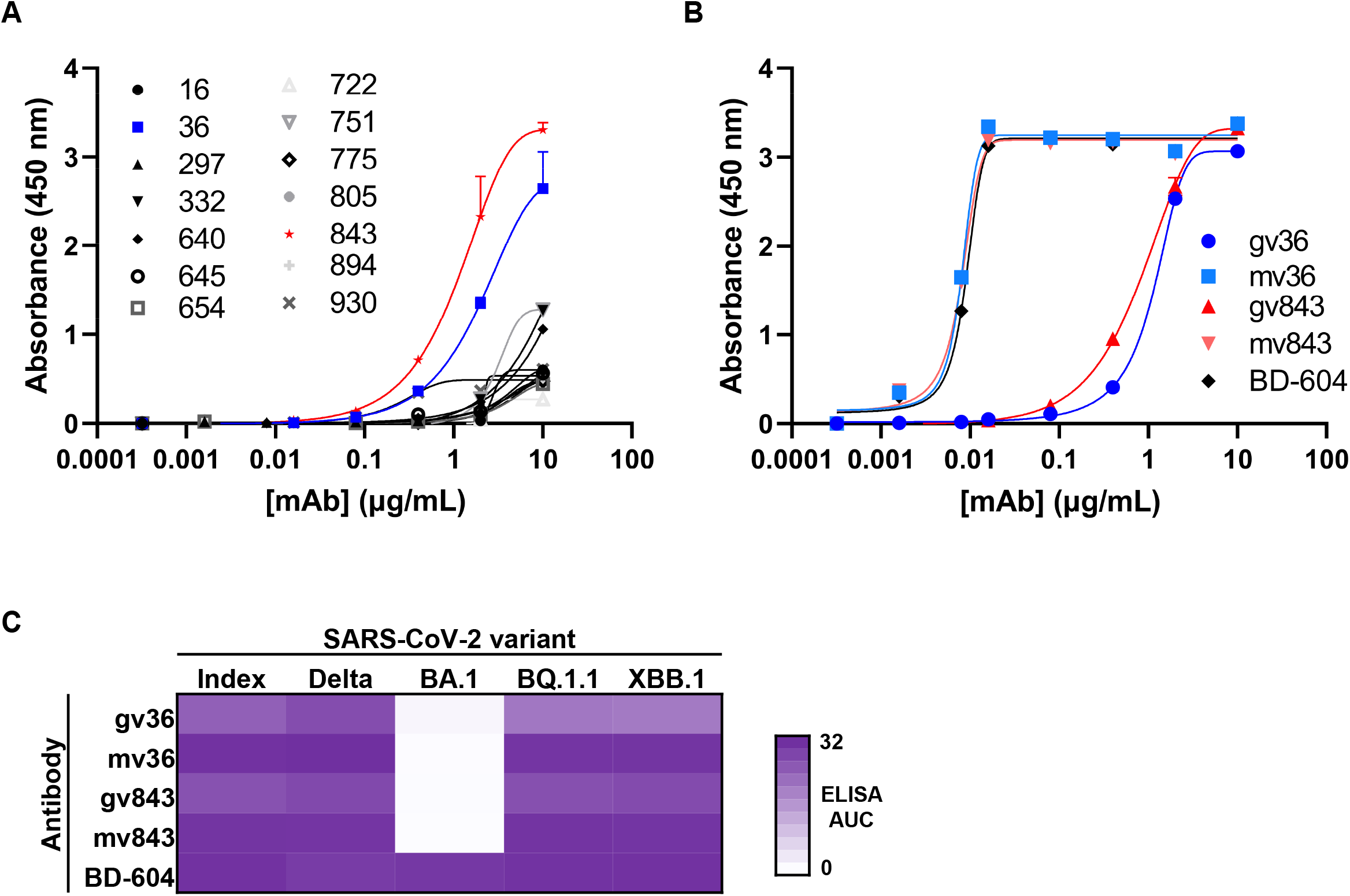
Binding activity of AI-generated antibodies against SARS-CoV-2 spike. (A) ELISA binding of AI-generated antibody candidates (symbols) to SARS-CoV-2 spike index variant. Data are mean + error of technical duplicates from a representative experiment repeated twice. (B) ELISA binding of germline and mature VH versions of antibodies 36 (blue) and 843 (red) to SARS-CoV-2 spike index variant. Known VH3-53 antibody BD-604 was used as a control. Data are mean + error of technical duplicates from a representative experiment repeated twice. (C) Heatmap of ELISA area under the binding curve (AUC) of germline and mature VH versions of antibodies 36 and 843 (rows) to different spike variants of SARS-CoV-2 virus. Known VH3-53 antibody BD-604 was used as a control.

Together, these results suggest that AI-based methods can successfully generate novel antibody CDRH3 sequences targeting a specific antigen of interest, and that predicted structural similarity to known antigen-specific antibodies can be used to down-select candidates among the initial pool of generated sequences. The specific pipeline that we applied here incorporates several variables, the optimization of which can further improve the success of the models – for example, the methods for modeling the structure of the AI-designed antibodies. Nevertheless, it is encouraging that multiple antigen-specific antibodies were identified through validating a small number of candidates, with a notable hit rate (∼15%). As generative antibody sequence algorithms and down-selection modeling approaches continue to be optimized, the efficiency and accuracy of generating antigen-specific antibodies through AI technologies will be expected to increase immensely, and to expand beyond CDR-only sequence generation and into truly de novo antibody design.

With traditional antibody discovery technologies, the targeted identification of antibodies against an antigen of interest requires complex experimental protocols ^18^, and is rarely capable of identifying antibodies against a specific epitope on the antigen target ^19^. In contrast, the computational pipeline described here can be applied to successfully and efficiently identify epitope-specific antibodies given an existing antibody-epitope template, which will be a valuable step forward for the antibody discovery field. The process of antibody generation in nature is highly complex, yet AI-based technologies appear capable of reproducing the end output based on the same building blocks but without explicit knowledge about the actual relationships and rules that underlie this process in nature. It is interesting to note that using a germline-level gene as a building block is sufficient for the generation of antibodies with reasonable binding to the target antigen. The observed notable increase in antigen binding when a small set of (5) mutations are introduced into the VH region of the antibodies suggests that combining AI-based antibody generation with established AI approaches for antibody affinity maturation (such as ^8^) can efficiently result in the design of high-affinity antibodies. Overall, our results point to the potential of AI technologies for de novo antibody design and will serve as the basis for further technology development that will enable the targeted and template-free generation of antigen-specific antibodies with high affinity, breadth of reactivity, and other properties of translational interest.

## MATERIALS AND METHODS

### De Novo Sequence Generation and Computational Analysis

The HC sequence of BD-604 was reverted to its V3-53/J6 germline sequence other than the CDRH3 region (i.e., from framework 1 to framework 3, as well as framework 4). The reverted sequence was used as a template for the de novo generation of CDRH3 sequences by using the IgLM^13^ infill function with chain token “[HEAVY]” and species token “[HUMAN]” and amino acid infill range corresponding to the CDRH3 region (ALA.96-VAL.106). The generated 1,000 CDRH3 sequences were encoded with the ANN4D^20^ encoding scheme and clustered using HDBCSAN^21^. The sequences were reordered in the resulting 12 clusters and pairwise sequence identity was calculated. The sequence identity between two sequences Seq1 and Seq2 was defined as 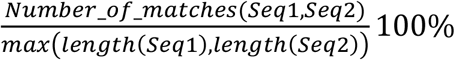. The structures of all the generated antibody HC (the de novo generated CDRH3 and the flanking V/J) were modeled by employing the NanoBodyBuilder2 function of ImmuneBuilder^15^. For structural similarity estimation, the unadjusted RMSD (calculated in PyMOL Open Source 2.5.0) was used.

### Antigen expression and purification

SARS-CoV-2 spike index, delta, BA.1, BQ.1.1, and XBB.1 variants were transiently expressed in Expi293F cells in FreeStyle F17 expression media (Thermo Fisher) (0.1% Pluronic Acid F-68 and 20% 4 mM L-glutamine) using the Expifectamine transfection reagents (Thermo Fisher Scientific). Cultures were suspended for 6 days with shaking at 8% CO_2_ saturation. At harvest, cultures were centrifuged at a minimum of 6000 rpm for 20 minutes before filtering supernatant with Nalgene Rapid Flow Disposable Filter Units with PES membrane (0.45 or 0.22 μm). Supernatant was passed over a 1 mL pre-packed StrepTrap XT column (Cytiva Life Sciences) equilibrated with binding buffer (100 mM Tris-HCl, 150 mM NaCl, 1 mM EDTA, pH 8.0). Following the binding phase, the column was washed with 15 mL of binding buffer before 10 mL of elution buffer (binding buffer supplemented with 2.5 mM desthiobiotin) was passed over the column to collect purified protein. Purified protein in elution buffer was concentrated before buffer exchanging into PBS, and then run on a Superose 6 Increase 10/300 GL on the AKTA FPLC system. Size exclusion peaks that corresponded to trimeric spike species were confirmed via SDS-PAGE of elution fractions. Fractions representative of pure trimeric spike were pooled and quantified using UV/vis spectroscopy. ELISA binding with known monoclonal antibodies was used to confirm antigenicity of spike proteins. Proteins were aliquoted, frozen, and kept at - 80°C until use.

### Antibody expression and purification

Antibody constructs were synthesized by Twist BioScience. Breifly, variable genes of each tested antibody were inserted into bi-cistronic plasmids encoding the constant regions for the heavy chain and either the kappa or lambda light chain. mAbs were expressed by transient transfection in Expi293F cells in FreeStyle F17 expression media (Thermo Fisher) (0.1% Pluronic Acid F-68 and 20% 4 mM L-glutamine) using the Expifectamine transfection reagents (Thermo Fisher Scientific). Cultured were suspended for 5 days with shaking at 8% CO_2_ saturation and 37°C. Five days post transfection, cultures were harvested and centrifuged at a minimum of 6000 rpm for 20 minute before filtration with Nalgene Rapid Flow Disposable Filter Units with PES membrane (0.45 or 0.22 μm). Filtered supernatant was run over PBS-equilibrated columns containing 1 ml of Protein A agarose resin. Following the binding phase, columns were washed with PBS and then purified antibodies were eluted with 5 mls of 100 mM Glycine HCl at pH 2.7 directly into 1 mL of 1 M Tris-HCl pH 8 and then buffer exchanged into PBS.

### Enzyme linked immunosorbent assay (ELISA)

Recombinant antigen was plated at a concentration of 2 μg/ml and left at 4°C overnight. The following day, plates were coated with 1% BSA in PBS supplemented with 0.05% Tween20 (PBS-T) for one hour at room temperature, Tested antibodies were diluted in 1% BSA in PBS-T, starting at 10 μg/ml and serially diluted five fold seven times for one hour incubation.

Secondary antibody, goat anti-human IgG conjugated to peroxidase, was used at 1:10,000 dilution in 1% BSA in PBS-T, for a one hour incubatino prior to development by adding TMB substrate to each well. TMB was incubated at room temperature for five minutes before adding 1 N sulfuric acid to quench the reaction. Plates were read at 450 nm. Plates were washed three times with PBS-T between all steps. ELISAs were performed in technical and biological duplicate.

## ACKNOWLEDGMENTS

This work was conducted, in part, using the resources of the Advanced Computing Center for Research and Education (ACCRE) at Vanderbilt University. For work described in this manuscript, I.S.G., T.M.M., A.A.A., and A.K.J. were supported in part by the G. Harold and Leila Y. Mathers Charitable Foundation (MF-2107-01851) and NIH R01AI175245 (to I.S.G.). A.A.A. was also supported in part by NIH T32AI112541. The funders had no role in the conceptualization or execution of any studies or drafting of the manuscript.

## DATA AVAILABILITY

Data in this manuscript is available upon request to the corresponding author, Ivelin Georgiev.

## References

1. Castelli, M.S., McGonigle, P. & Hornby, P.J. The pharmacology and therapeutic applications of monoclonal antibodies. Pharmacology Research & Perspectives 7, e00535 (2019).

2. Kinch, M.S., Kraft, Z. & Schwartz, T. Monoclonal antibodies: Trends in therapeutic success and commercial focus. Drug Discovery Today 28, 103415 (2023).

3. Shiakolas, A.R. et al. Efficient discovery of SARS-CoV-2-neutralizing antibodies via B cell receptor sequencing and ligand blocking. Nat Biotechnol (2022).

4. Akbar, R. et al. Progress and challenges for the machine learning-based design of fit-for-purpose monoclonal antibodies. mAbs 14, 2008790 (2022).

5. Wilman, W. et al. Machine-designed biotherapeutics: opportunities, feasibility and advantages of deep learning in computational antibody discovery. Briefings in Bioinformatics 23 (2022).

6. Ding, W., Nakai, K. & Gong, H. Protein design via deep learning. Briefings in Bioinformatics 23 (2022).

7. Wang, H. et al. Scientific discovery in the age of artificial intelligence. Nature 620, 47–60 (2023).

8. Hie, B.L. et al. Efficient evolution of human antibodies from general protein language models. Nature Biotechnology (2023).

9. Parkinson, J., Hard, R. & Wang, W. The RESP AI model accelerates the identification of tight-binding antibodies. Nature communications 14, 454 (2023).

10. Wang, Y., Yuan, M., Lv, H., Peng, J., Wilson, I.A. & Wu, N.C. A large-scale systematic survey reveals recurring molecular features of public antibody responses to SARS-CoV-2. Immunity 55, 1105–1117. e1104 (2022).

11. Zhang, Q. et al. Potent and protective IGHV3-53/3-66 public antibodies and their shared escape mutant on the spike of SARS-CoV-2. Nature communications 12, 4210 (2021).

12. Ferruz, N. & Höcker, B. Controllable protein design with language models. Nature Machine Intelligence 4, 521–532 (2022).

13. Shuai, R.W., Ruffolo, J.A. & Gray, J.J. IgLM: Infilling language modeling for antibody sequence design. Cell Syst (2023).

14. Olsen, T.H., Moal, I.H. & Deane, C.M. AbLang: an antibody language model for completing antibody sequences. Bioinformatics Advances 2 (2022).

15. Abanades, B., Wong, W.K., Boyles, F., Georges, G., Bujotzek, A. & Deane, C.M. ImmuneBuilder: Deep-Learning models for predicting the structures of immune proteins. Communications Biology 6, 575 (2023).

16. Du, S. et al. Structurally Resolved SARS-CoV-2 Antibody Shows High Efficacy in Severely Infected Hamsters and Provides a Potent Cocktail Pairing Strategy. Cell 183, 1013–1023.e1013 (2020).

17. He, Q. et al. An updated atlas of antibody evasion by SARS-CoV-2 Omicron sub-variants including BQ.1.1 and XBB. Cell Rep Med 4, 100991 (2023).

18. Setliff, I. et al. High-throughput mapping of B-cell receptor sequences to antigen specificity. Cell 179, 1636–1646 (2019).

19. Walker, L.M. et al. High-Throughput B Cell Epitope Determination by Next-Generation Sequencing. Front Immunol (2022).

20. Meiler, J., Müller, M., Zeidler, A. & Schmäschke, F. Generation and evaluation of dimension-reduced amino acid parameter representations by artificial neural networks. Molecular modeling annual 7, 360–369 (2001).

21. Campello, R.J.G.B., Moulavi, D. & Sander, J. 160–172 (Springer Berlin Heidelberg, Berlin, Heidelberg; 2013).

